# Patterns of gene content and co-occurrence constrain the evolutionary path toward animal association in CPR bacteria

**DOI:** 10.1101/2021.03.03.433784

**Authors:** Alexander L. Jaffe, Christine He, Ray Keren, Luis E. Valentin-Alvarado, Patrick Munk, Keith Bouma-Gregson, Ibrahim F. Farag, Yuki Amano, Rohan Sachdeva, Patrick T. West, Jillian F. Banfield

## Abstract

Candidate Phyla Radiation (CPR) bacteria are small, likely episymbiotic organisms found across Earth’s ecosystems. Despite their prevalence, the distribution of CPR lineages across habitats and the genomic signatures of transitions amongst these habitats remain unclear. Here, we expand the genome inventory for Absconditabacteria (SR1), Gracilibacteria, and Saccharibacteria (TM7), CPR bacteria known to occur in both animal-associated and environmental microbiomes, and investigate variation in gene content with habitat of origin. By overlaying phylogeny with habitat information, we show that bacteria from these three lineages have undergone multiple transitions from environmental habitats into animal microbiomes. Based on co-occurrence analyses of hundreds of metagenomes, we extend the prior suggestion that certain Saccharibacteria have broad bacterial host ranges and constrain possible host relationships for Absconditabacteria and Gracilibacteria. Full-proteome analyses show that animal-associated Saccharibacteria have smaller gene repertoires than their environmental counterparts and are enriched in numerous protein families, including those likely functioning in amino acid metabolism, phage defense, and detoxification of peroxide. In contrast, some freshwater Saccharibacteria encode a putative rhodopsin. For protein families exhibiting the clearest patterns of differential habitat distribution, we compared protein and species phylogenies to estimate the incidence of lateral gene transfer and genomic loss occurring over the species tree. These analyses suggest that habitat transitions were likely not accompanied by large transfer or loss events, but rather were associated with continuous proteome remodeling. Thus, we speculate that CPR habitat transitions were driven largely by availability of suitable host taxa, and were reinforced by acquisition and loss of some capacities.

**IMPORTANCE:** Studying the genetic differences between related microorganisms from different environment types can indicate factors associated with their movement among habitats. This is particularly interesting for bacteria from the Candidate Phyla Radiation because their minimal metabolic capabilities require symbiotic associations with microbial hosts. We found that shifts of Absconditabacteria, Gracilibacteria, and Saccharibacteria between environmental ecosystems and mammalian mouths/guts probably did not involve major episodes of gene gain and loss; rather, gradual genomic change likely followed habitat migration. The results inform our understanding of how little-known microorganisms establish in the human microbiota where they may ultimately impact health.

## INTRODUCTION

The Candidate Phyla Radiation (CPR) is a phylogenetically diverse clade of bacteria characterized by reduced metabolisms, potentially episymbiotic lifestyles, and ultrasmall cells. While the first high-quality CPR genomes were primarily from groundwater, sediment, and wastewater (1–3), subsequently genomes have been recovered from diverse environmental and animal-associated habitats, including humans. Intriguingly, from dozens of major CPR lineages, only three – Candidatus Absconditabacteria (formerly SR1), Gracilibacteria (formerly BD1-5 and GN02), and Saccharibacteria (formerly TM7) – are consistently associated with animal oral cavities and digestive tracts (4). The Saccharibacteria are perhaps the most deeply studied of all CPR lineages to date, likely due to their widespread presence in human oral microbiomes and association with disease states such as gingivitis and periodontitis (5, 6). On the other hand, Absconditabacteria and Gracilibacteria remain deeply undersampled, potentially due to their rarity in microbial communities or their use of an alternative genetic code that may confound some gene content analyses (1, 7, 8).

Absconditabacteria, Gracilibacteria, and Saccharibacteria are predicted to be obligate fermenters, dependent on other microorganisms (hosts) for components such as lipids, nucleic acids, and many amino acids (1, 3). Despite a generally reduced metabolic platform, CPR bacteria display substantial variation in their genetic capacities, even within lineages (9, 10). For example, some Gracilibacteria lack essentially all genes of the glycolysis and pentose-phosphate pathways and the TCA cycle (11). In contrast to many CPR, soil-associated Saccharibacteria encode numerous genes related to oxygen metabolism (12, 13). Pangenome analyses have shown genetic evidence for niche partitioning among Saccharibacteria from the same body site (14). However, the lack of comprehensive genomic sampling of these three CPR lineages across habitats, particularly from environmental biomes, has left unclear the full extent to which CPR gene inventories vary with habitat type, and, relatedly, the extent to which changes in metabolic capacities might have been reshaped during periods of environmental transition. Of particular interest is whether rapid gene acquisitions (e.g., via lateral gene transfer) or losses enabled habitat switches, or if these changes occurred gradually following habitat change. The availability of suitable hosts may also drive the colonization of new environments by CPR bacteria (14). While there has been significant progress in characterizing the relationship between Saccharibacteria and Actinobacteria in the oral habitat (15–17), other CPR-host relationships remain unclear. Elucidation of environmental transitions among CPR lineages will require both thorough analysis of functional repertoires as well as a more comprehensive understanding of associations with other microorganisms. Here, we improve existing sampling of CPR genomes and their surrounding communities to examine patterns of distribution, abundance, and gene content in different microbiome types. We also make use of whole-community co-occurrence patterns to shed light on the potential host range of the CPR bacteria in their associated ecosystems. In combination, our analyses shed light on the frequency of habitat shifts in three CPR lineages and the evolutionary processes likely underlying them.

## RESULTS

### Environmental diversity, phylogenetic relationships, and abundance patterns

We gathered an environmentally comprehensive set of Absconditabacteria, Gracilibacteria, and Saccharibacteria by querying multiple databases for genomes assembled in previous studies and assembling new genomes from several additional metagenomic data sources (Table S1, Materials and Methods) (1–4, 7, 12–15, 18–82). Quality filtration of this curated genome set at ≥ 70% completeness and ≤ 10% contamination and subsequent de-replication at 99% average nucleotide identity (ANI) yielded a non-redundant set of 389 genomes for downstream analysis (Table S1). Absconditabacteria and Gracilibacteria were less frequently sampled relative to Saccharibacteria, comprising only ∼7.5% and ∼10.8% of the total genome set, respectively. All three lineages were distributed across a broad range of microbiomes, encompassing various environmental habitats (freshwater, marine, soil, engineered, plant-associated, hypersaline) as well as multiple animal-associated microbiomes (oral and gut) (Fig. 1). Unlike animal-associated Gracilibacteria and Absconditabacteria genomes, which were recovered only from human and animal oral samples, animal-associated Saccharibacteria were found in both oral and gut samples.

**Fig 1.**
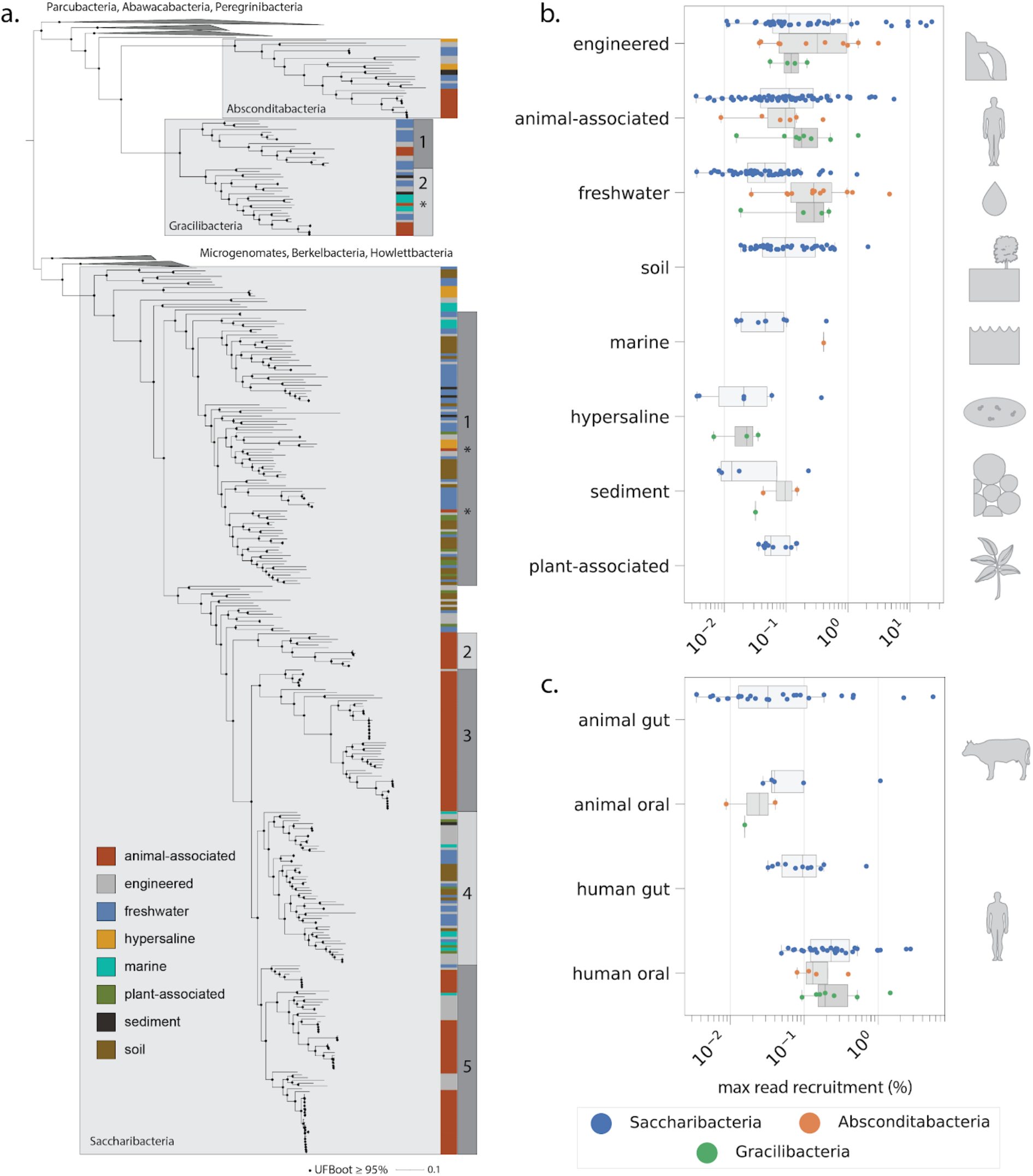
Phylogenetic and environmental patterns for the Absconditabacteria, Gracilibacteria, and Saccharibacteria. **a)** Maximum-likelihood tree based on 16 concatenated ribosomal proteins (1976 amino acids, LG+R10 model). Scale bar represents the average number of substitutions per site. Asterisks indicate phylogenetic position of a subset of organisms derived from dolphin mouth metagenomes. Percentage of reads per metagenomic sample mapping to individual genomes across **b)** environments and **c)** body sites of humans and animals.

We extracted 16 syntenic, phylogenetically informative ribosomal proteins from each genome to construct a CPR species tree and evaluate how habitat of origin maps onto phylogeny.

Sequences from related CPR bacteria were used as outgroups for tree construction (Materials and Methods). The resolved topology supports monophyly of all three lineages and a sibling relationship between the two alternatively coded lineages, Absconditabacteria and Gracilibacteria (Figure 1a, File S1), consistent with previous findings for the CPR (10). For the Absconditabacteria, a single clade of organisms derived from animal-associated microbiomes was deeply nested within genomes from the environment. On the other hand, Gracilibacteria clearly formed two major lineages (GRA 1-2), each with a small subclade comprised of animal-associated genomes. For Saccharibacteria, deeply-rooting lineages were also almost exclusively of environmental origin (soil, water, sediment) and animal-associated genomes were strongly clustered into at least three independent subclades (Fig. 1a). Two of these three subclades were exclusively composed of animal-associated sequences whereas one (SAC 5), was a mixture of animal-associated, wastewater (potentially of human origin) and a few aquatic sequences. Intriguingly, for both Saccharibacteria and Gracilibacteria, a subset of organisms from the dolphin mouth (22) did not affiliate with those from terrestrial mammals/humans and instead fell within marine/environmental clades (indicated by asterisks in Fig. 1). In primarily environmental clades (SAC 1 and 4), genomes from soil, freshwater, engineered, and halophilic environments were phylogenetically interspersed, suggesting comparatively wide global distributions for these lineages. One exception to this pattern were two clades representing distinct hypersaline environments – a hypersaline lake and salt crust (65, 70).

We used read mapping to assess the abundance of Absconditabacteria, Gracilibacteria, and Saccharibacteria genomes in the samples from which they were originally reconstructed. Generally, these CPR bacteria are not dominant members of microbial communities (<1% of reads). However, they were relatively abundant in some engineered, animal-associated, and freshwater environments (Fig. 1b). In rare cases, CPR taxa comprised >10% of reads (Fig. 1b), and in a bioreactor (engineered) reached a maximum of ∼22% of reads. Gracilibacteria and Absconditabacteria attained comparable read recruitment to Saccharibacteria and were particularly abundant in some groundwater and animal-associated habitats. In contrast to Saccharibacteria, Gracilibacteria and Absconditabacteria have so far only been minimally detected in soil and plant-associated microbiomes. We also compared abundance patterns across animal body sites. As expected based on extensive prior work (5, 16, 44), Saccharibacteria exhibited highest read recruitment in the human oral microbiome. However, these bacteria can also comprise a significant fraction of the microbial community in exceptional gut/oral microbiomes from cows, pigs, and dolphins (Fig. 1c), in one case approaching 5% of sequenced reads (Table S2). When detected, Saccharibacteria in the human gut were relatively rare, comprising a median of ∼0.1% of reads across samples.

### Patterns of co-occurrence constrain CPR host range across environments

Despite recent progress made in experimentally identifying bacterial host range for oral Saccharibacteria, little is known about associations in other habitats. Abundance pattern correlations can be informative regarding associations involving obligate symbionts and their microbial hosts (37, 55); however, such analyses often rely on highly resolved time-series for statistical confidence. Here, we instead examine patterns of co-occurrence within samples to probe potential relationships between CPR bacteria and their microbial hosts. Given recent experimental evidence demonstrating the association of multiple Saccharibacteria strains with various Actinobacteria in the human oral microbiome (15–17, 44, 83), we predicted that Actinobacteria may be common hosts of Saccharibacteria in microbiomes other than the mouth and asked to what extent co-occurrence data supported this relationship.

We first identified all ribosomal protein S3 (rps3) sequences from Actinobacteria and Saccharibacteria in the source metagenomes probed in this study for relative abundance patterns (Fig. 1bc). Rps3 sequences from all samples were clustered into ‘species groups’ (Materials and Methods). We observed that species groups from Actinobacteria and Saccharibacteria frequently co-occurred in the soil and plant-associated microbiomes as well as several hypersaline microbiomes (Fig. 2a). On the other hand, co-occurrence of the two lineages was less frequent in engineered and freshwater environments relative to other environments. Surprisingly, only ∼78% of animal-associated samples containing Saccharibacteria also contained Actinobacteria at abundances high enough to be detected (Fig. 2a). Assemblies with well-sampled Saccharibacteria yet no detectible Actinobacteria could suggest that Saccharibacteria have alternative hosts in these samples or are able to (at least periodically) live independently.

**Fig 2.**
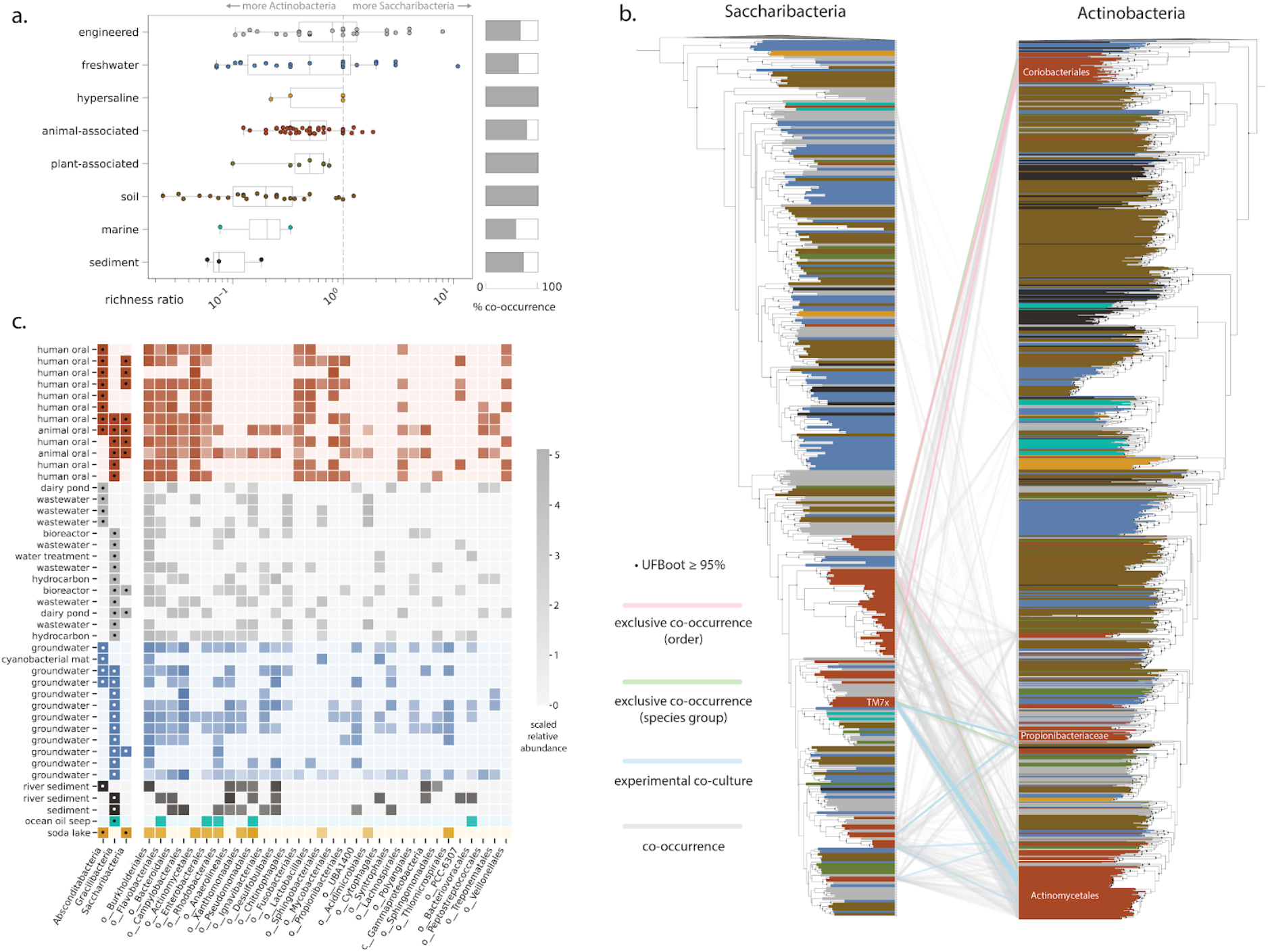
Patterns of co-occurrence between CPR and potential host lineages across environments. **a)** Relative richness ratio, describing the ratio of distinct Saccharibacteria species groups to Actinobacteria species groups, for each sample and overall co-occurrence percentage across habitat categories. **b)** Maximum-likelihood trees for Saccharibacteria and Actinobacteria based on ribosomal protein S3 sequences extracted from all source metagenomes. Co-occurrence patterns are shown only for species groups derived from animal-associated metagenomes. **c)** Community composition for metagenomic samples containing Absconditabacteria and Gracilibacteria. Cells with dots indicate only presence, whereas those without dots convey quantitative relative abundance information. Only potential host lineages present in 8 or more samples are shown.

For samples where both Saccharibacteria and Actinobacteria marker genes were detectable, we computed a ‘relative richness’ metric describing the ratio of distinct Saccharibacteria species groups to Actinobacteria species groups. In most animal-associated microbiomes, Actinobacteria were more species rich (lower richness ratios), as expected if individual Saccharibacteria can associate with multiple hosts (Fig. 2a). Greater species richness of Actinobacteria compared to Saccharibacteria was also observed for many plant-associated, soil, engineered, and freshwater microbiomes. However, some engineered freshwater samples had richness ratios equal to (equal richness) or greater than 1 (i.e., Saccharibacteria more species rich) (Fig. 2a). Specifically, we observed that several metagenomes from engineered and freshwater environments contained anywhere from 1-11 Saccharibacteria species but only one detectable Actinobacteria species (Table S4). Thus, if Actinobacteria serve as hosts for Saccharibacteria in these habitats, there may be both exclusive associations and associations linking multiple Saccharibacteria species with a single Actinobacteria host species.

We next tested for more specific possible associations in the animal microbiome, reasoning that if Actinobacteria are common hosts for Saccharibacteria, then exclusive co-occurrence of a particular Saccharibacteria species with singular Actinobacteria species within a sample might suggest an interaction *in vivo*. We mapped all pairs of Saccharibacteria and Actinobacteria species that co-occurred within a single sample onto the trees constructed from the rpS3 sequences (Fig. 2b), including 22 Saccharibacteria-Actinobacteria pairs reported in previous experimental studies (Table S3). In three additional cases, we found that individual metagenomic samples contained only one assembled Saccharibacteria species group and one Actinobacteria species group (“exclusive co-occurrence - species group”, Fig. 2b). Two of these cases involved Actinobacteria from the order Actinomycetales, from which multiple Saccharibacteria hosts have already been identified. We also noted exclusive species-level co-occurrence of a Saccharibacteria species group from the human gut and an Actinobacteria species group from the order Coriobacterales (Table S4). In an additional seven cases, one Saccharibacteria species group occurred with multiple Actinobacteria species groups of the same order-level classification based on rps3 gene profiling (“exclusive co-occurrence - order”, Fig. 2b). Five of the seven instances involved pairs of Saccharibacteria and Coriobacterales from termite and swine gut metagenomes. Thus, unlike in human oral environments, Coriobacterales may serve as hosts for Saccharibacteria in gut environments of multiple animal species. More generally, we also observed that Saccharibacteria from the same phylogenetic clade had predicted relationships to phylogenetically unrelated Actinobacteria (Fig. 2b), consistent with previous experimental observations for individual species (44).

Compared to Saccharibacteria, host relationships for Gracilibacteria and Absconditabacteria have received little attention. There are preliminary indications that Absconditabacteria may associate with members of the Fusobacteria or Firmicutes in the oral microbiome (44) or the gammaproteobacterium *Halochromatium* in certain salt lakes (84). We thus explored co-occurrence patterns in microbial communities containing Absconditabacteria and Gracilibacteria, attempting to further constrain possible host taxa. In animal and human-associated microbiomes, bacteria from several lineages, including Fusobacteria (Fig. 2c), were relatively abundant in nearly all samples that contained Absconditabacteria. Members of the Chitinophagales, Pseudomonadales, and Acidimicrobiales were detected in high abundance in three wastewater samples from similar treatment plants (40) and one dairy pond sample containing Absconditabacteria. No clear patterns of potential host co-occurrence were observed for Gracilibacteria, with the exception of the proteobacterial order Campylobacterales, which co-occurred in 8 of 10 groundwater samples where Gracilibacteria were found (Fig. 2c). Across all environments, only members of the order Burkholderiales (a large order of Gammaproteobacteria) consistently co-occurred with Gracilibacteria.

Among the least complex communities that contained Absconditabacteria were cyanobacterial mats from a California river network, where dominant cyanobacterial taxa accounted for ∼60-98% relative abundance (32). To complement the above co-occurrence analyses, we re-analyzed 22 published metagenomes representing spatially separated mats and discovered that Absconditabacteria were detectable in 12 of them at varying degrees of coverage (0.12-37x). As noted previously, also present in the mats were members of the phyla Bacteroidetes, Betaproteobacteria, and Verrucomicrobia (32). Correlation of read coverage profiles across mats provided moderate support for the association of Absconditabacteria and Bacteroidetes. Specifically, many of the strongest species-level correlations, including five of the top ten, involved Bacteroidetes (Table S5).

### Gene content of Absconditabacteria, Gracilibacteria, and Saccharibacteria

We next examined how gene content of these CPR lineages varied across environments. We first compared the predicted proteome size of these bacteria across habitats, taking into consideration differing degrees of genome completeness. This analysis revealed that genomes from soil and the rhizosphere (plant-associated) have on average larger predicted proteomes relative to those from animal-associated environments (Fig. 3b). Saccharibacteria from hypersaline environments appear to have the smallest predicted proteomes, although the limited number of high-quality genomes in this category currently limits a firm conclusion. We observed some evidence for variance in predicted proteome sizes among Absconditabacteria and Gracilibacteria, including potentially smaller predicted proteomes among animal-associated Gracilibacteria (Fig. S1). Additional high quality genomes will be required to confirm this trend.

**Fig 3.**
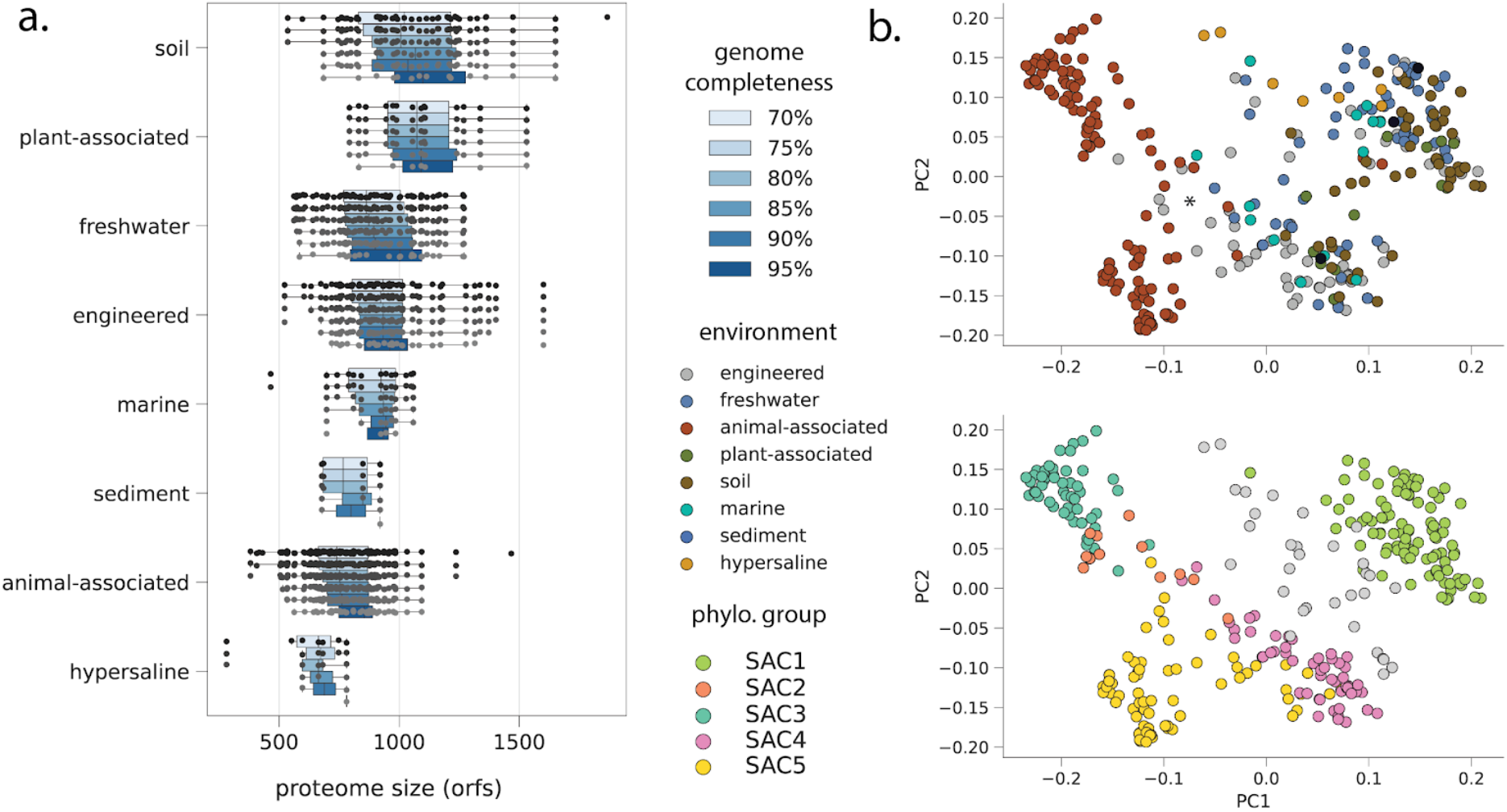
Proteome characteristics for Saccharibacteria. **a)** Predicted proteome size (open reading frame count) at increasing genome completeness thresholds. **b)** Overall proteome similarity among Saccharibacteria from different environmental categories (top panel) and phylogenetic clades (bottom panel). PCoAs were computed from presence/absence profiles of all protein clusters with 5 or more member sequences. The primary (PC1) and secondary (PC2) axis of variation explained 12% and 8% of variance, respectively.

To examine overall proteome similarity as a function of habitat type, we employed a recently developed protein-clustering approach that is agnostic to functional annotation (9) (Table S6, Materials and Methods). Among Saccharibacteria, principal coordinates analysis (PCoA) of presence/absence profiles for all protein families with 5 or more members yielded a primary axis of variation (∼12% variance explained) that distinguished animal-associated Saccharibacteria from environmental or plant-associated ones and a secondary axis (∼8 % variance explained) that distinguished between phylogenetic clades (SAC1-3 vs 4-5). We did not observe strong clustering of Saccharibacteria by specific environmental biome, consistent with the interspersed nature of their phylogenetic relationships (Fig. 1a, 3b). Notably, several SAC5 genomes from wastewater have protein family contents that are intermediate between those of animal-associated Saccharibacteria and Saccharibacteria from the large environmental clade (indicated by an asterisk in Fig. 3b). This finding may indicate selection within the engineered environments for variants introduced from human waste. PCoAs of predicted proteome content among Absconditabacteria and Gracilibacteria generally showed that, with the exception of dolphin-derived genomes, animal-associated lineages are also distinct from their relatives from environmental biomes (Fig. S2). Overall, our results indicate that the CPR lineages examined here have predicted proteomes whose content and size vary substantially with their environment. This is particularly evident for animal-associated Saccharibacteria, which are notably dissimilar in their protein family content compared to environmental counterparts.

To further examine the distinctions evident in the PCoA analysis, we arrayed presence/absence information for each protein family and hierarchically clustered them based on their distribution patterns across all three CPR phyla. This strategy allowed us to explore specific protein family distributions and to test for groups of co-occurring protein families (modules) that are common to bacteria from a single lineage or are shared by most bacteria from one or more CPR lineages. We first observed one large module that is generally conserved across all genomes. This module is comprised of families for essential cellular functions such as transcription, translation, cell division, and basic energy generating mechanisms (Fig. 4a, “core”).

**Fig 4.**
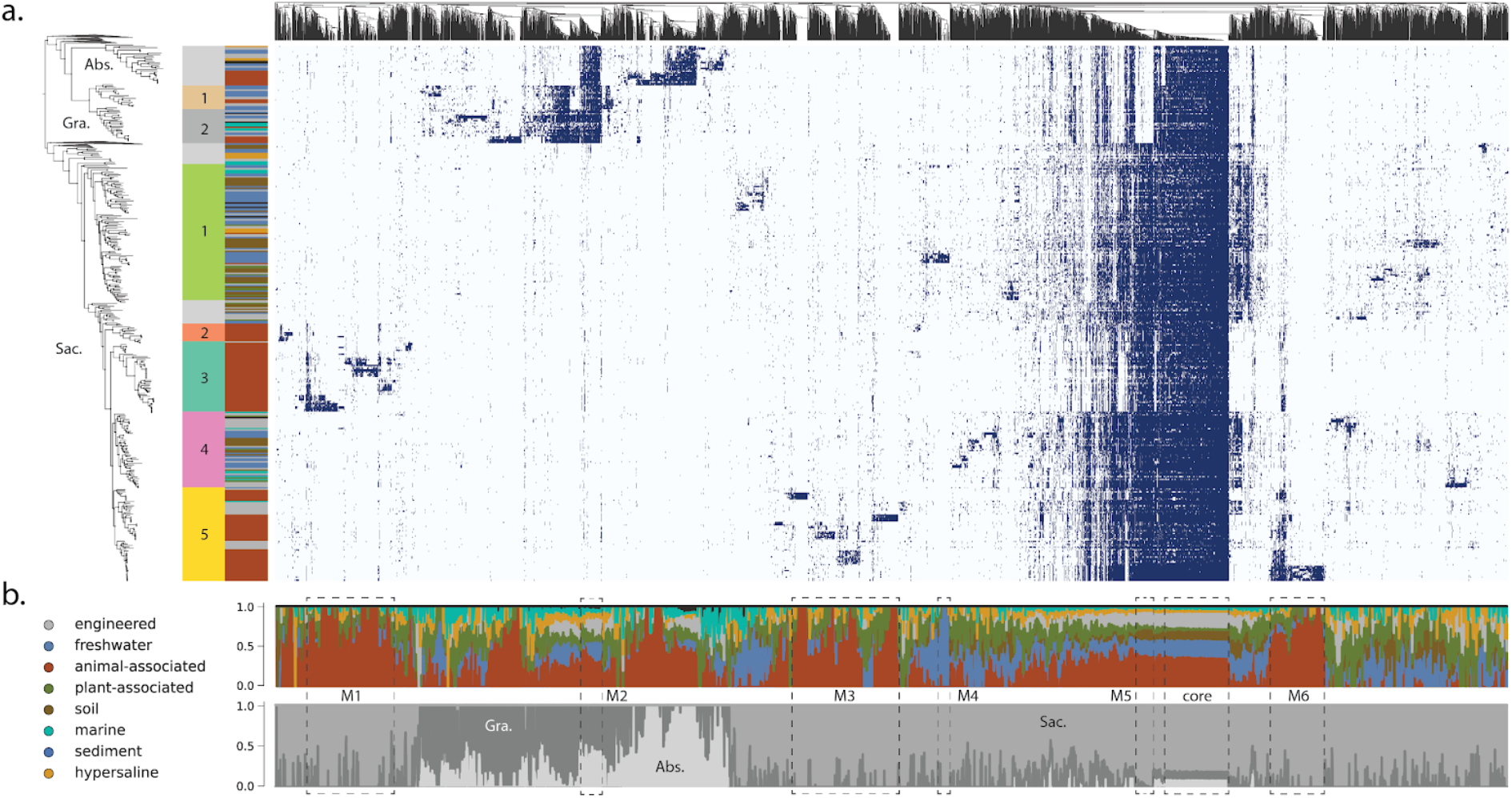
Phylogenetic and environmental distribution of protein families recovered among CPR. **a)** Presence/absence profiles for protein families with 5 or more members, with shaded cells indicating presence, and light cells indicating absence. Heatmap columns represent protein families, hierarchically clustered by similarity in distribution across the genome set. Rows correspond to genomes, ordered by their phylogenetic position in the species tree (left). Abbreviations: Abs., Absconditabacteria; Gra., Gracilibacteria; Sac., Saccharibacteria. **b)** Percentage of genomes encoding individual protein families that belonged to broad habitat groups (top panel) or taxonomic groups (bottom panel). Modules of protein families indicated in the text are represented by dotted lines (M1-6 and ‘core’).

The protein family analysis also revealed multiple modules specific to Gracilibacteria, Absconditabacteria, and modules shared by both lineages but not present in Saccharibacteria, paralleling their phylogenetic relationships (Fig. 1a, 4a). Of the ∼70 families shared only by Gracilibacteria and Absconditabacteria (M2, Fig. 4a), nearly half had no KEGG annotation at the thresholds employed. One family shared by these phyla but not in Saccharibacteria is the ribosomal protein L9, which supports prior findings on the composition of Saccharibacteria ribosomes (29). The remaining families also include two that were fairly confidently annotated as the DNA mismatch repair proteins, MutS and MutL (fam01378 and fam00753), nicking endonucleases involved in correction of errors made during replication (85) (Table S6). Despite the generally wide conservation of these proteins among Bacteria, we saw no evidence for the presence of either enzyme in Saccharibacteria, suggesting that aspects of DNA repair may vary in this group relative to other CPR. We recovered a module of approximately 60 proteins highly conserved among the Saccharibacteria and only rarely encoded in the other lineages (M5, Fig. 4a). This module contained several protein families confidently annotated as core components of glycolysis and the pentose phosphate pathway, including three enzymes present in almost all CPR (10): glyceraldehyde 3-phosphate dehydrogenase, (GAPDH) triosephosphate isomerase (TIM), and phosphoglycerate kinase (PGK). These results indicate that Gracilibacteria and Absconditabacteria may have extremely patchy, if not entirely lacking, components of core carbon metabolism, even when a high-quality genome set is considered.

For all three lineages of CPR, we also observed numerous small modules with narrow distributions. To test whether these modules represent functions differentially distributed among organisms from different habitats, we computed ratios describing the incidence of each protein family in one habitat compared to all others (Materials and Methods). Enriched families were defined as those with ratios ≥ 5, whereas depleted families were defined as those that were encoded by <10% of genomes in a given habitat, but ≥ 50% of genomes from other habitats. To account for the fact that small families might appear to be differentially distributed due to chance alone, we also stipulated that comparisons be statistically significant (p ≤ 0.05, two-sided Fisher’s exact test corrected for multiple comparisons).

Using this approach, we identified 926 families that were either enriched (n=872) or depleted (n=54) in genomes from one or more broad habitat groups. We identified 45 families enriched in Absconditabacteria from animal-associated environments relative to those from environmental biomes. The majority of these families were either poorly functionally characterized or entirely without a functional annotation at the thresholds employed. Similarly, families enriched in animal-associated Gracilibacteria relative to environmental counterparts were primarily unannotated; among those families with confident annotations was a family likely encoding a phosphate:Na+ symporter (fam04488) and a putative membrane protein (fam06579). Intriguingly, 6 families were co-enriched in both animal-associated Gracilibacteria and Absconditabacteria, suggesting that these sibling lineages might have acquired or retained a small complement of genes that are important in adaptation to animal habitats or their associated bacteria.

Animal-associated Saccharibacteria, on the other hand, encoded 417 unique families that were exclusive or highly enriched relative to those from other habitats. Enriched families largely fell into three major groups (M1, M3, M6; Fig. 4), and the large majority of them, particularly among modules with narrow, lineage-specific distributions, were without functional annotations. However, our analysis also revealed some protein families with broader distributions across multiple clades of animal-associated Saccharibacteria (Fig. 4). Here, among families with functional annotations, we found several apparently involved in the transport of amino acids and dicarboxylates that were highly enriched (ratios ranging from 10 to 112.9) in the majority of animal-associated Saccharibacteria (52-58% of genomes across clades) (Table S7). Two of these families, corresponding to a putative amino acid transport permease and substrate-binding protein (fam00393 and fam11477, respectively) were co-located in some genomes along with a ATP-binding protein (subset of fam00001), suggesting that they may function together to uptake amino acids. We also recovered several other functions that were previously predicted to be enriched based on analysis of a smaller set of animal-associated Saccharibacteria (4), including phosphoglycerate mutase, glycogen phosphorylase, and a uracil-DNA glycosylase (ratio 8.3-33.5). Lastly, we found that the CRISPR-associated protein cas9 was moderately enriched among animal-associated genomes (ratio ∼5 among 33 genomes), consistent with the suggestion that these Saccharibacteria likely acquired their viral defense systems after colonizing animals (Table S7) (4).

We identified multiple families that are either enriched or depleted in animal-associated Saccharibacteria that were functionally related to oxidative stress (Table S7). Among enriched families, one (fam00662) set was mostly annotated with low confidence as rubrerythrin, a family of iron containing proteins generally involved in detoxification of peroxide (86). Member sequences of this family were present in over a third of animal-associated Saccharibacteria and were highly enriched relative to environmental genomes (fold-enrichment ratio of 36.2), suggesting that acquisition may have conferred an adaptive benefit in the gut and/or oral cavity. In contrast, we also observed that animal-associated Saccharibacteria were significantly depleted in another family confidently annotated as a Fe-Mn family superoxide dismutase (fam01569) and likely involved in radical detoxification. Animal-associated lineages were also strongly depleted for the genes comprising the cytochrome o ubiquinol oxidase operon (fam00281, fam00112, fam01347, fam00624, and fam10494), with very few, if any, animal-associated genomes and more than 50% of environmental genomes encoding each of the five genes. This operon has been previously suggested to confer an advantage in aerophilic environments like soil through detoxification (3) or use of oxygen (12, 13).

Among genomes from environmental biomes, we identified a module of approximately 100 protein families, also primarily without functional annotation, that were associated with a subclade of Saccharibacteria recently reconstructed from metagenomes of freshwater lakes and glacier ice (M4, Fig. 4) (61, 87). Notably, among the most widespread families in this module was one in which sequences were annotated as bacteriorhodopsin with low confidence (fam11249). Further analysis indicated that these sequences fall within the bacterial/archaeal Type 1 rhodopsin clade and contain both the retinal-binding lysine associated with light sensitivity and a DTS motif (Fig. S3), suggesting that they may function as proton pumps (88, 89). Distinct rhodopsin sequences were also recovered in the genomes of environmental Absconditabacteria (NDQ motif) and Gracilibacteria (DTE motif), although they were not statistically enriched (Fig. S3). Genomes of soil-associated Saccharibacteria were enriched for about 130 protein families largely without strong functional annotations (Fig. 3). Despite their small proteome sizes, Saccharibacteria from hypersaline environments were only statistically depleted in about 15 families at the thresholds employed here. Sequence files for all protein families are provided in the Supplementary Materials (File S2).

### Evolutionary processes shaping proteome evolution

The observation that some differentially distributed traits among CPR were apparently lineage specific, whereas others were more widespread, motivated us to examine the relative contributions of gene transfer and loss to proteome evolution. To do so, we first inferred unrooted, maximum-likelihood phylogenies for the sequences in each protein family that was differentially distributed, then compared these phylogenies to the previously reconstructed species tree (Materials and Methods). For each family, the likelihood of transfer and loss events on each branch of the species tree were then estimated using a probabilistic framework that takes into consideration genome incompleteness, variable rates of transfer and loss, and uncertainty in gene tree reconstruction (90, 91). The results of this analysis reveal relatively few instances of originations, defined as lateral transfer from outside the three lineages of CPR or *de novo evolution* (‘originations’, Fig. 5). In the Absconditabacteria and Gracilibacteria, gene-species tree reconciliation revealed that small modules of families of mostly hypothetical proteins were acquired near the base of animal-associated clades (O1-O2, Fig. 5). On the other hand, in Saccharibacteria, originations were primarily associated with shallower subclades of animal-associated (and in one case, freshwater) genomes (O3-O6, Fig. 5). These findings generally corresponded with the distribution of small, highly enriched modules of largely hypothetical proteins (Fig. 4) and suggest that the distribution of these modules is best explained by lineage-specific acquisition events of relatively few genes at one time, rather than large acquisition events at deeper nodes. Intriguingly, one subclade of animal-associated Saccharibacteria had the highest incidence of originations of all groups in our analysis (O6), suggesting that these genomes may be phylogenetic ‘hotspots’ for transfer.

**Fig 5.**
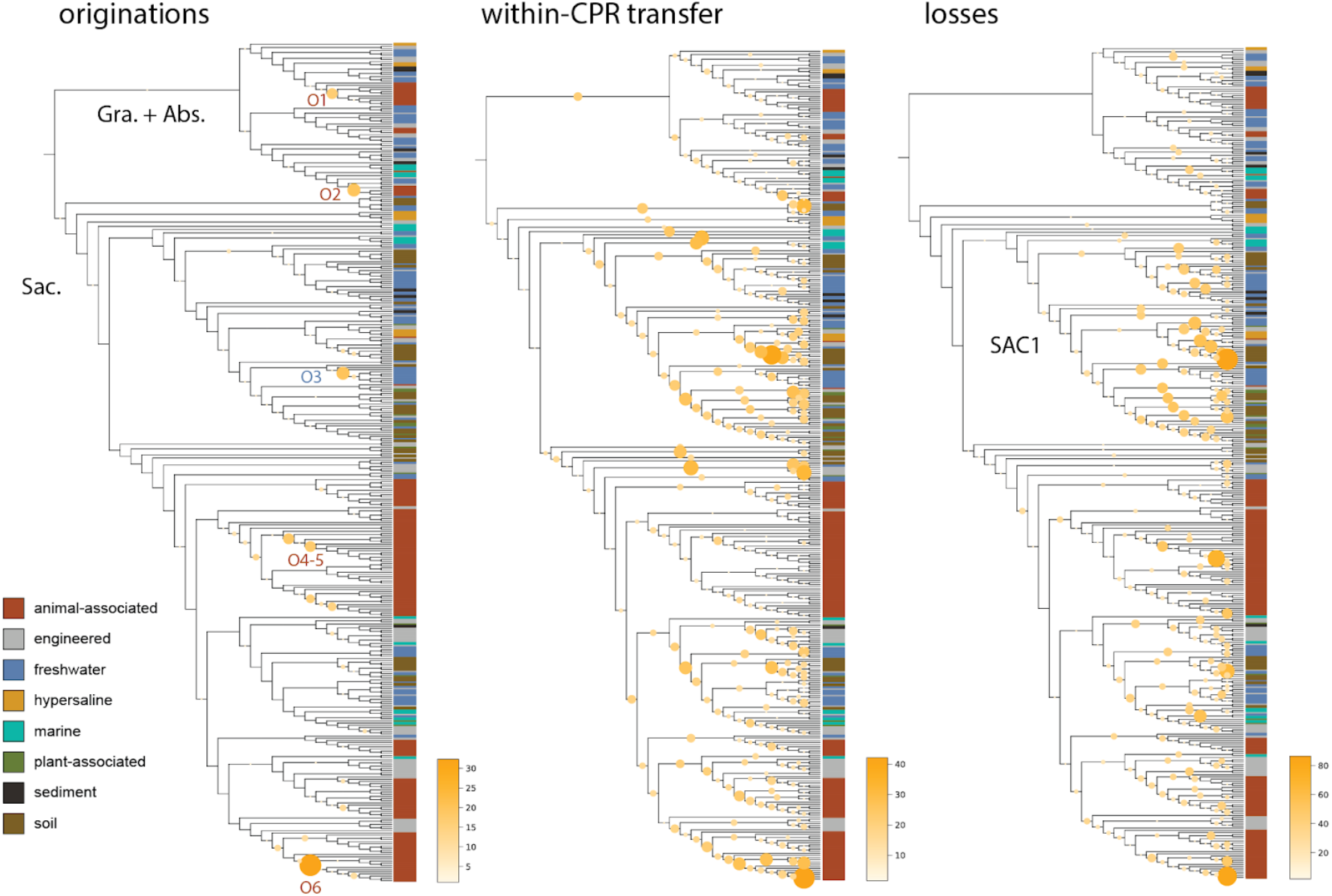
Evolutionary processes shaping proteome evolution in three lineages of CPR bacteria. Each panel displays the species tree from Fig. 1 in cladogram format. The size and color of circles mapped onto interior branches represent the cumulative number of **a)** originations (defined as either lateral transfer from outside the lineages examined here, or *de novo* evolution) **b)** transfer among the three CPR lineages included here and **c)** genomic losses predicted to occur on that branch for all 902 differentially distributed families where gene-species tree reconciliation was possible. Abbreviations: Abs., Absconditabacteria; Gra., Gracilibacteria; Sac., Saccharibacteria. SAC1 indicates a monophyletic clade of Saccharibacteria referenced in the text.

While origination events were relatively infrequent in all three CPR lineages, instances of within-CPR transfer and loss were very frequent and dispersed across most interior branches of the tree (Fig. 5). Notably, we detected sporadic losses across internal branches, which is inconsistent with a major gene loss event at the time of adaptation to animal-associated habitats. Surprisingly, we noticed that genomes of non-animal associated Saccharibacteria, particularly those from the SAC1 clade, displayed substantial patterns of loss despite their relatively large proteome sizes. Thus, losses in these environmental lineages were possibly balanced by lateral transfer events over the course of evolution.

## DISCUSSION

Here, we expand sampling of genomes from the Absconditabacteria, Gracilibacteria, and Saccharibacteria, particularly from environmental biomes. The basal positioning of environmental clades in phylogenetic reconstructions provides strong support for the hypothesis that these lineages originated in the environment (Fig. 1a), and potentially migrated into humans and terrestrial animals via consumption of groundwater (4, 41). Unlike the Absconditabacteria, which appear to have transitioned only once into animal oral cavities and guts, our phylogenetic evidence suggests that Gracilibacteria and Saccharibacteria have undergone multiple transitions into the animal microbiome in unique phylogenetic clades. This includes Saccharibacteria and Gracilibacteria from the dolphin mouth that appear to have colonized these marine mammals from a distinct source compared to those that colonized the oral environments of terrestrial animals (Fig. 1). In other clades, phylogenetically interspersed environmental and oral/gut Saccharibacteria could reflect either independent migrations into the animal environment or lineage-specific reversion back to environmental niches (SAC2-5, Fig. 1).

Currently, the mechanisms that enable environmental transition among CPR are unknown. Several observations, including that CPR host-pairs may be taxonomically distinct between oral and gut habitats, raise the question of whether habitat transitions among CPR involve co-migration with their hosts or the acquisition of new hosts. The finding that single CPR species co-occur with a single Actinobacteria species, or several closely related ones, in multiple animal-associated metagenomes contributes further evidence that these associations can be flexible and phylogenetically diverse rather than highly evolutionarily conserved (44). Supporting this, some laboratory strains of oral Saccharibacteria can adapt to new hosts after periods of living independently (92). The lack of evidence for lateral gene transfer between experimentally profiled pairs (4) also suggests that some CPR-host pairs may have established fairly recently.

Host associations for Absconditabacteria, Gracilibacteria, and environmental Saccharibacteria are currently unknown. Previously, changes in abundance over a sample series from a bioreactor system treating thiocyanate was used to suggest that *Microbacterium ginsengisoli* may serve as a host for a co-occurring Saccharibacteria (55). One Absconditabacteria lineage (*Vampirococcus*) has been predicted to have a host from the Gammaproteobacteria (84) and one Gracilibacteria was suggested to have a *Colwellia* host based on a shared repeat protein motif (11). Given the scant information on hand about possible hosts for these CPR, especially for Absconditabacteria and Gracilibacteria, the patterns of co-occurrence we report for specific organisms provide starting points for host identification via targeted co-isolation.

To evaluate to what extent changes in gene content are associated with habitat transition, we first established core gene sets indicated that overall proteome size and content differed between environmental and animal-associated Saccharibacteria, and to some extent Gracilibacteria. Despite overall smaller proteome size, we identified a large number of protein families that were highly enriched among animal-associated CPR from all three lineages. The most striking capacities involve amino acid transport, oxidative stress tolerance, and viral defense, which may enable use of habitat-specific resources or tolerance of habitat-specific stressors. These findings complement previous suggestions that prophages are enriched in animal-associated Saccharibacteria relative to environmental counterparts (14).

Only three lineages of CPR (of potentially dozens) have been consistently recovered in the animal-associated microbiome. Given the enormous diversity of CPR bacteria in drinking water (41), there has likely been ample opportunity for various taxa to disperse into the mouths of terrestrial animals; however, establishment and persistence of these bacteria may have been limited by the absence of a suitable host in oral and gut environments. Thus, we predict that other CPR bacteria - including those from the large Microgenomates and Parcubacteria lineages - have hosts that are infrequent or transient members of the animal microbiome, or have insufficient ability to ‘adapt’ to new hosts upon contact. For example, formation of new associations may be limited by the specificity of pili involved in host interaction or proteins involved in attachment (14, 41, 93).

It is also interesting to compare processes of habitat transition in CPR with those proposed for other bacteria and for archaea. Our results suggest that Saccharibacteria (and potentially Gracilibacteria) from the human/animal microbiome have smaller genome sizes than related, deeper-branching lineages of environmental origin. This pattern is also apparent for other, free-living groups adapted to the animal microbiome from the environment, like the Elusimicrobia (94) and intracellular symbionts of insects (95). Furthermore, in contrast to findings for Elusimicrobia, where host-associated lineages have common patterns of loss of metabolic capacities compared to relatives from non-host environments (94), patterns of gene loss in animal-associated CPR appear to be heterogeneous and lineage-specific. One possibility is that gene loss in CPR is primarily modulated by strong dependence on host bacteria, whose capacities may vary substantially, rather than by adaptation to the relatively stable, nutrient-rich animal habitat that likely shaped evolution of some non-CPR bacteria.

Changes in gene content could enable, facilitate, or follow habitat transitions. Our evolutionary reconstructions revealed that habitat-specific differences in gene content are more likely the product of combinations of intra-CPR transfer and loss rather than major acquisition events at time of lineage divergence. Thus, modules enriched in specific lineages were probably acquired via lateral transfer after habitat transition, suggesting that proteome remodeling has been continuous in CPR over evolutionary time. As such, the processes shaping CPR lineage evolution share both similarities and differences with those predicted for other microbes, including Haloarchaeota (96) and ammonia-oxidizing lineages of Thaumarchaeota (90, 97), where both large lateral transfer events and gradual patterns of gene loss, gain, and duplication worked together to shape major habitat transitions.

## CONCLUSION

Overall, our findings highlight factors associated with habitat transitions in three CPR lineages that occur in both human/animal and environmental microbiomes. We expand the evidence for niche-based differences in protein content (4, 14) and identify a large set of protein families that could guide future studies of CPR symbiosis. Furthermore, patterns of co-occurrence may inform experiments aiming to co-cultivate CPR and their hosts. Our analyses point to a history of continuous genome remodeling accompanying transition into human/animal habitats, rather than rapid gene gain/loss around the time of habitat switches. Thus, habitat transitions in CPR may have been primarily driven by the availability of suitable hosts and reinforced by acquisition and/or loss of genetic capacities. These processes may be distinct from those shaping transitions in other bacteria and archaea that are not obligate symbionts of other microorganisms.

## MATERIALS AND METHODS

### Genome database preparation and curation

To compile an environmentally comprehensive set of genomes from the selected CPR lineages, we first queried four genomic information databases - GTDB (https://gtdb.ecogenomic.org/), NCBI assembly (https://www.ncbi.nlm.nih.gov/assembly), PATRIC (https://www.patricbrc.org/), and IMG (https://img.jgi.doe.gov/) - for records corresponding to the Absconditabacteria, Gracilibacteria, and Saccharibacteria genomes. Genomes gathered from these databases were combined with those drawn from several recent publications as well as genomes newly binned from metagenomic samples of sulfidic springs, an advanced treatment system for potable reuse of wastewater, human saliva, cyanobacterial mats, fecal material from primates, baboons, pigs, goats, cattle, and rhinoceros, several deep subsurface aquifers, dairy-impacted groundwater and associated enrichments, multiple bioreactors, soil, and sediment (Table S1). Assembly, annotation, and binning procedures followed those from Anantharaman et al. 2016. In some cases, manual binning of the alternatively coded Absconditabacteria was aided by a strategy in which a known Absconditabacteria gene was blasted against predicted metagenome scaffolds to find ‘seed’ scaffolds, whose coverage and GC profile were used to probe remaining scaffolds for those with similar characteristics. For newly binned genomes, genes were predicted for scaffolds > 1 kb using prodigal (meta mode) and annotated using usearch against the KEGG, UniProt, and UniRef100 databases. Bins were ‘polished’ by removing potentially contaminating scaffolds with phylogenetic profiles that deviated from consensus taxonomy at the phylum level. One genome was further manually curated to remove scaffolding errors identified by read mapping, following the procedures outlined in (98).

We removed exact redundancy from the combined genome set by identifying identical genome records and selecting one representative for downstream analyses. We then computed contamination and completeness for the genome set using a set of 43 marker genes sensitive to described lineage-specific losses in the CPR (29, 31) using the custom workflow in checkm (99). Results were used to secondarily filter the genome set to those with ≥ 70% of the 43 marker genes present and ≤ 10% of marker genes duplicated. The resulting genomes were then de-replicated at 99% ANI using dRep (-sa 0.99 -comp 70 -con 10) (100), yielding a set of 389 non-redundant genomes. Existing metadata were used to assign both “broad” and “narrow” habitat of origin for each non-redundant genome. The “engineered” habitat category was defined to include human-made or industrial systems like wastewater treatment, bioreactors, and water impacted by farming/mining. Curated metadata, along with accession/source information for each genome in the final set, is available in Table S1. All newly binned genomes are available through Zenodo (Data and Software Availability).

### Functional annotation and phylogenomics

We predicted genes for each genome using prodigal (“single” mode), adjusting the translation table (*-g 25*) for Gracilibacteria and Absconditabacteria, which are known to utilize an alternative genetic code (7, 8). Predicted proteins were concatenated and functionally annotated using kofamscan (101). Results with an e-value ≤ 1e-6 were retained and subsequently filtered to yield the highest scoring hit for each individual protein.

To create a species tree for the CPR groups of interest, functional annotations from kofamscan were queried for 16 syntenic ribosomal proteins (rp16). Marker genes were combined with those from a set of representative sequences of major, phylogenetically proximal CPR lineages (10). Sequences corresponding to each ribosomal protein were separately aligned with MAFFT and subsequently trimmed for phylogenetically informative regions using BMGE (*-m BLOSUM30*) (102). We then concatenated individual protein alignments, retaining only genomes for which at least 8 of 16 syntenic ribosomal proteins were present. A maximum-likelihood tree was then inferred for the concatenated rp16 (1976 amino acids) set using ultrafast bootstrap and IQTREE’s extended Free-Rate model selection (*-m MFP -st AA -bb 1000*) (103). The maximum likelihood tree is available as File S1. The tree and associated metadata were visualized in iTol (104) where well-supported, monophyletic subclades were manually identified within Gracilibacteria and Saccharibacteria for use in downstream analysis.

### Abundance analysis

To assess the global abundance of Absconditabacteria, Gracilibacteria, and Saccharibacteria, we manually compiled the original read data associated with each genome in the analysis set, where available. We included only those genomes from short-read, shotgun metagenomics of microbial entire communities (genomes derived from single cell experiments, stable isotope probing experiments, “mini” metagenomes, long-read sequencing experiments, and co-cultures were excluded). For each sequencing experiment, we downloaded the corresponding raw reads and, where appropriate, filtered out animal-associated reads by mapping to the host genome using bbduk (*qhdist*=*1*). Sequencing experiments downloaded from the NCBI SRA database were sub-sampled to the average number of reads across all compiled experiments (∼36 million) using seqtk (*sample -s 7*) if the starting read pair count exceeded 100 million. We then removed Illumina adapters and other contaminants from the remaining reads and further quality trimmed them using Sickle. The filtered read set was then mapped against all genomes assembled (or co-assembled) from it using bowtie2 (default parameters). For mappings with a non-zero number of read alignments, relative abundance of each genome was calculated by counting the number of stringently mapped (≥99% identity) using coverm (*--min-read-percent-identity 0*.*99*) and dividing by the total number of reads in the quality-filtered read set. Genomes were considered present in a sample if at least 10% of sequence length was covered by reads. In cases where genomes were derived from co-assemblies of multiple sequencing experiments, we computed the abundance for each sample individually and then selected the one with the highest value as a ‘representative’ sample for downstream analyses.

### Co-occurrence analyses

Each representative sample was then probed for co-occurrence patterns of CPR and potential host lineages. To account for across-study differences in binning procedures, quality-filtered read sets were re-assembled using megahit (*--min-contig-len 1000*) and subsequently analyzed using graftm (105) with a ribosomal protein S3 (rps3) gpackage custom built from GTDB (release 05-RS95). Recovered rps3 protein sequences in each sample were clustered to form ‘species groups’ at 99% identity using usearch cluster_fast (*-sort length -id 0*.*99*). For all samples with >0 marker hits, we then performed three downstream analyses to examine patterns of co-occurrence for various taxa. First, we counted the number of unique species groups in each sample taxonomically annotated as Saccharibacteria (*“c Saccharimonadia”*) and Actinobacteria (“*p Actinobacteriota”*), dividing the former by the latter to compute a species ‘richness ratio’ for each sample (where *p Actinobacteria* did not equal 0). Second, to examine the co-occurrence of Saccharibacteria and Actinobacteria within a phylogenetic framework, we inferred maximum likelihood trees for the set of rps3 marker genes recovered across samples. Species group sequences were clustered *across* samples to further reduce redundancy using usearch (as described above) and were combined with rps3 sequences drawn from a taxonomically balanced set of bacterial reference genomes (10) as an outgroup. Saccharibacteria and Actinobacteria sequence sets were then aligned, trimmed, and used to build trees as described above for the 16 ribosomal protein tree, with the exception of using trimal (*-gt 0*.*1*) (106) instead of BMGE. Species groups that co-occurred in one or more metagenomic samples were then linked. If a given Saccharibacteria species group exclusively co-occurred with an Actinobacteria species group in *at least* one sample, or Actinobacteria species groups belonging to the same order level in *all* samples, those linkages were labelled. Finally, experimental co-cultures of Saccharibacteria and Actinobacteria from previous studies were mapped onto the trees. To do this, we compiled a list of strain pairs and their corresponding genome assemblies (Table S3) and then used graftm to extract rps3 sequences from corresponding genome assemblies downloaded from NCBI. We then matched these rps3 sequences to their closest previously defined species group using blastp (*-evalue 1e-3-max_target_seqs 10 -num_threads 16 -sorthits 3 -outfmt 6*), prioritizing hits with the highest bitscore and alignment length. Reference rps3 sequences with no match at ≥99% identity and ≥95% coverage among the species groups were inserted separately into the tree. We then labelled all experimental pairs of species in the linkage diagram.

Third, we profiled a subset of 43 metagenomes containing Gracilibacteria and Absconditabacteria for overall community composition. For each sample, we extracted all contigs bearing ribosomal S3 proteins and mapped the corresponding quality filtered read set to them using bowtie2. Mean coverage for each contig was then computed using coverm (*contig --min-read-percent-identity* .*99*) and a minimum covered fraction of 0.10 was again employed. Relative coverage for each order level lineage (as predicted by graftm) was computed by summing the mean coverage values for all rps3-bearing contigs belonging to that lineage.

Where species groups did not have order-level taxonomic predictions, the lowest available rank was used. Finally, relative coverage values were scaled by first dividing by the lowest relative coverage observed across samples and then taking the base-10 log. For the re-analysis of 22 cyanobacterial mat metagenomes (32), the same approach was taken, and coverage profiles for rps3-bearing scaffolds were correlated using the pearsonr function in the scipy.stats package.

### Proteome size, content, enrichment

We subjected all predicted proteins from the genome set to a two-part, de novo protein clustering pipeline recently applied to CPR genomes, in which proteins are first clustered into “subfamilies” and highly similar/overlapping subfamilies are merged using and HMM-HMM comparison approach (--coverage 0.70) (9) (https://github.com/raphael-upmc/proteinClustering-Pipeline). For each protein cluster, we recorded the most common KEGG annotation among its member sequences and the percent of sequences bearing this annotation (e.g. 69% of sequences in fam00095 were matched with K00852).

We then performed three subsequent analyses to describe broad proteome features of included CPR. First, we computed proteome size across habitats, defined as the number of predicted ORFs per genome when considering genomes at increasing thresholds of completeness in single copy gene inventories (75%, 80%, 85%, etc.). Second, we examined similarity between proteomes by generating a matrix describing the presence/absence patterns of protein families with 5 or more member sequences. We then used this matrix to compute distance metrics between each genome based on protein content using the ecopy package in Python (method=‘jaccard’, transform=‘1’) and performing a principal coordinates analysis (PCoA) using the skbio package. The first two axes of variation were retained for visualization alongside environmental and phylogenetic metadata. Finally, we used the clustermap function in seaborn (*metric*=*‘jaccard’, method*=*‘average’*) to hierarchically cluster the protein families based on their distribution patterns and plot these patterns across the genome set. For each protein family, we also computed the proportion of genomes encoding at least one member sequence that belonged to each of the three CPR lineages and each broad environmental category (Fig. 4b) (see custom code linked in Data and Software Availability).

We next identified protein families that were differentially distributed among genomes from broad environmental categories. For each protein family, we divided the fraction of genomes from a given habitat (‘in-group’) encoding the family by the same fraction for genomes from all other habitats (‘out-group’). In cases where no ‘out-group’ genome encoded a member protein, the protein family was simply noted as ‘exclusive’ to the ‘in-group’ habitat. In all cases, we calculated the Fisher’s exact statistic using the *fisher_exact* function in scipy.stats (*alternative*=*‘two-sided’*). To account for discrepancies in genome sampling among lineages, we determined ratios and corresponding statistical significance values separately for each lineage. All statistical comparisons for a given lineage were corrected for false discovery rate using the *multipletests* function in statsmodels.stats.multitest (*method*=*“fdr_bh”*). Finally, we selected families that were predicted to be enriched or depleted in particular habitats. We considered enriched families to be those with ratios ≥ 5, and depleted families as those that were encoded in 10% or fewer of genomes from a given habitat but present in 50% or more of genomes outside that environmental category. Retaining only those comparisons with corrected Fisher’s statistics at 0.05 or below resulted in a set of 926 unique, differentially distributed protein families for downstream analysis.

### Analysis of putative rhodopsins

Protein sequences from the CPR (fam11249) were combined with a set of reference protein sequences spanning Type 1 bacterial/archaeal rhodopsin and heliorhodopsin (107). Sequences were then aligned using MAFFT (*--auto*) and a tree was inferred using FastTreeMP (default parameters). Alignment columns with 95% or more gaps were trimmed manually in Geneious for the purposes of visualization. Transmembrane domains were identified by BLASTp searches (https://blast.ncbi.nlm.nih.gov/Blast.cgi) and conserved residues were defined by manual comparison with an annotated alignment of previously published reference sequences (108).

### Processes driving protein family evolution

To examine the evolutionary processes shaping the differentially-distributed protein families, we next subjected each family to an automated gene-species tree reconciliation workflow adapted from (90). Briefly, for each family, truncated sequences (defined as those with lengths less than 2 standard deviations from the family mean) were removed and the remaining sequences aligned with MAFFT (*--retree 2*). Resulting alignments were then trimmed using trimal (*-gt 0*.*1*) and used to infer maximum-likelihood phylogenetic trees using IQTree with 1000 ultrafast bootstrap replicates (*-bnni -m TEST -st AA -bb 1000 -nt AUTO*). We removed reference sequences from the inferred species tree and rooted it on the branch separating Saccharibacteria from the monophyletic clade containing Gracilibacteria and Absconditabacteria. A random sample of 100 bootstrap replicates were then used to probabilistically reconcile each protein family with the pruned species tree using the ALE package (*ALE_undated*) (91). Estimates of missing gene fraction were derived from the checkM genome completeness estimates described above. We then calculated the total number of originations (horizontal gene transfer from non-CPR, or *de novo* gene formation), within-CPR horizontal transfers, and losses over each non-terminal branch and mapped branch-wise counts for each event to a species-tree cladogram in iTol (104).

## SUPPLEMENTARY MATERIAL

All supplementary figures, tables, and files are available through Zenodo (https://doi.org/10.5281/zenodo.4586041).

**Fig. S1**. Proteome size as a function of genome completeness and habitat of origin for Absconditabacteria and Gracilibacteria.

**Fig. S2**. Principal coordinates analysis based on all protein families with 5 or more members among **ab)** all lineages, **c)** Absconditabacteria, and **d)** Gracilibacteria.

**Fig. S3**. Phylogenetic relationships and trimmed protein alignment among Type 1 rhodopsins and CPR rhodopsin homologs. CPR sequences from the Absconditabacteria (NDQ motif), Gracilibacteria (DTE motif), and Saccharibacteria (DTS motif) are highlighted in the tree. Sequence conservation at each aligned site, the location of bacteriorhodopsin (BR) site 85, 89, 96, and the location of retinal-binding lysine (Schiff Base linkage) are also indicated.

**File S1**. Absconditabacteria, Gracilibacteria, and Saccharibacteria species tree based on 16 syntenic ribosomal proteins (newick format).

**File S2**. FASTA-formatted files for 3,787 protein families with 5 or more member sequences.

**Table S1**. Characteristics of genomes used in this study, including environmental metadata and accession information.

**Table S2**. Read accession and mapping information for genomes included in the global abundance analysis.

**Table S3**. Metadata and ribosomal protein S3 (rpS3) information for experimentally validated Saccharibacteria-Actinobacteria pairs. Only strains with publicly available reference genomes and detectable rpS3 sequences are listed.

**Table S4**. Pairs of Saccharibacteria-Actinobacteria species groups with exclusive co-occurrence at the species group or order level in the included metagenomic samples.

**Table S5**. Top 25 coverage correlations between Absconditabacteria and other organisms (as denoted by representative rpS3-bearing scaffolds) across 22 cyanobacterial mat metagenomes.

**Table S6**. Characteristics of the 3,787 protein families with 5 or more member sequences.

**Table S7**. Characteristics of protein families that are statistically enriched/depleted across habitat categories.

## Acknowledgments

We thank Shufei Lei, Lily Law, Alex Crits-Christoph, Tom Williams, Oded Béjà, Adair Borges, Raphaël Méheust, Alison Sharrar, Alexa Nicolas, Jett Liu, and Simonetta Gribaldo for informatics support, helpful discussions, and comments on the manuscript. We thank Alex Thomas, Ariel Amadio, Mircea Podar, Ramunas Stepanauskas, Connor Skennerton, Stefano Campanaro, Cédric Laczny, Paul Wilmes, Clara Chan, Scott E. Miller, Lauren C. Kennedy, Rose S. Kantor, Kara L. Nelson, Lauren Lui, Maliheh Mehrshad, Chris Greening, Mads Albertsen, and Sari Peura for permission to use genomic data that were unpublished at the time of writing.

Christine He was funded by a Camille & Henry Dreyfus Environmental Chemistry Postdoctoral Fellowship. Patrick Munk was supported by the Danish Veterinary and Food Administration and The Novo Nordisk Foundation (NNF16OC0021856). JAEA was funded by the Ministry of Economy, Trade and Industry of Japan, as “The Project for Validating Near-field System Assessment Methodology in Geological Disposal System”. Keith Bouma-Gregson and cyanobacterial mat sample collection was supported by the National Science Foundation’s Eel River Critical Zone Observatory [EAR-1331940], Department of Energy grant [DOE-SC10010566], NSF Division of Environmental Biology [1656009], and US EPA STAR Fellowship [91767101-0]. Rohan Sachdeva and thiocyanate reactor genome construction was supported by a grant from the National Science Foundation (USA) to JFB (EAR-1349278). We also thank the Innovative Genomics Institute at UC Berkeley.

## Author’s contributions

A.L.J and J.F.B. compiled the dataset, performed genome curation and analysis, developed the project, and wrote the manuscript. C.H., R.K., L.E.V.A., P.M., K.B.G., I.F.F., Y.A., R.S., and P.T.W. generated data for the study and provided comments on the manuscript.

## Data and software availability

All accession information for the genomes analyzed in this study are listed in Supplementary Table 1. Genomes as well as custom code for the described analyses are also available on GitHub: https://github.com/alexanderjaffe/cpr-crossenv.

